# Using Deep Learning to Count Monarch Butterflies in Dense Clusters

**DOI:** 10.1101/2021.07.23.453502

**Authors:** Shruti Patel, Amogh Kulkarni, Ayan Mukhopadhyay, Karuna Gujar, Jaap de Roode

**Author notes:** Equal Contribution (Shruti Patel, Amogh Kulkarni).

## Abstract

Monarch butterflies display one of the most fascinating migration patterns of all species, traveling over 3000 miles from their North American breeding grounds to reach overwintering sites in Central Mexico. Recent studies have suggested that monarchs have experienced an alarming decline in population size due to a combination of deforestation, loss of native milkweed and nectaring plants, and climate change. An issue that conservation efforts face is the lack of principled mechanisms to accurately estimate and count the population size of monarchs. This difficulty occurs due to their small size and existence in dense overwintering clusters in forests. We create an open-source tool to aid conservationists estimate the count of monarch butterflies from images automatically. To the best of our knowledge, our approach, based on deep convolutional neural networks, is the first automated application that can count small insects like monarch butterflies in dense clusters. We demonstrate that our approach achieves high accuracy in counting the number of butterflies even in the presence of occlusion. We also release an open-source dataset containing high resolution images of monarch butterflies along with human annotations for each butterfly’s position. Our open-source implementation can be readily used by scientists to estimate monarch numbers in overwintering clusters and could also be adapted for use in other clustering species.

## 1 Introduction

Every year, billions of animals undertake long-distance migrations to leave deteriorating habitats, escape parasites and predators, and benefit from available resources in multiple regions [Dingle, 2014, Alerstam et al., 2003, Altizer et al., 2011]. Migrations across continents are one of the most fascinating phenomena that exist in nature. One of the most spectacular migrations is undertaken by the monarch butterfly in North America, with up to hundreds of millions of monarchs flying up to 3,000 miles to reach their overwintering grounds in Central Mexico [Reppert and Roode, 2018, Urquhart and Urquhart, 1978, Brower, 1995]. Monarch caterpillars are specialist feeders on milkweed plants, and breed between March and October across their 4.5 million km^2^ breeding range spanning the United States and southern Canada [Flockhart et al., 2013]. Upon milkweed deterioration, diminishing daylight, and decreasing temperature, monarchs embark on their journey south to overwinter in Oyamel forests in the states of Mexico and Michoacán, where temperatures are mild enough to prevent freezing, but low enough to prevent high activity and premature depletion of lipid reserves. In early spring, monarchs mate and fly north to recolonize their breeding grounds over 2-4 successive generations [Flockhart et al., 2013].

There are many reasons why monarch butterflies have become an active area of research. While these butterflies are deep-rooted in many Native American folktales and traditions in North America [Agrawal, 2017], they became the topic of much scientific study due to their particularly fascinating migration patterns and warning coloration [Brower, 1988]. More recently, monarchs have garnered much attention from the research community and public because of conservation concerns [Vidal and Rendón-Salinas, 2014]. Indeed, studies have suggested that the monarch population has undergone a drastic decline over the past three decades, with some estimates of a greater than 80% decline between a high in the winter of 1996–1997 and a low in the winter of 2014–2015 [Thogmartin et al., 2017b]. Moreover, some estimates indicate that the butterflies face quasi-extinction of 11–57% in the next 20 years [Semmens et al., 2016]. A major issue with these studies is that they are entirely based on estimates of occupied hectares in the overwintering sites. Thus, instead of actual population censuses, monarch population size has been estimated based on the number of hectares occupied by overwintering monarch clusters. Rough estimates of numbers of monarchs per hectare vary widely, and have been cited to range from 7 and 61 million [Thogmartin et al., 2017a]. With such diverging estimates, it is difficult to judge the actual monarch population size and how it has changed over time. Indeed, because of these uncertainties, and because monarch censuses during other parts of the annual life cycle do not indicate declines, some authors have suggested that monarchs may not actually be declining [Mawdsley et al., 2020, Davis, 2020].

Arguably, the most important step in conserving a species is to monitor its population. While this step might seem relatively straightforward for many species around the globe, it is particularly challenging to design principled methodologies to count and track monarch butterflies. While many citizen science monitoring projects are based on standardized efforts of monarch sightings during the breeding and migration season, it is the overwintering population that could provide the most accurate population estimates. This is because all monarchs gather in relatively small areas, and are relatively immobile during their overwintering phase. However, this approach introduces challenges. First, monarch butterflies stay in extremely dense clusters consisting of thousands of individuals. Second, they rest on trees at heights that are difficult to deal with from the perspective of manual counting. Finally, the clusters are oftentimes located in dense forests that are difficult to survey manually. The lack of counting methodologies for individual butterflies has resulted in estimating population size based on the area occupied by the butterflies. In principle, it should be feasible to combine this area method with counts of individual monarchs, counting the number of butterflies in a small sample of clusters to estimate the total number of butterflies. While such an approach is scalable, a high level of accuracy of the counting procedure in the sub-sample would be required. In this paper, we present an automated approach to count monarch butterflies (and possibly other species) in dense clusters.

### Contributions

**1)** We discuss a deep learning architecture that can count monarch butterflies from images by converting the images into a density map, and then learning a function that models the conditional distribution of the density map given the original image. **2)** We release an open source dataset of annotated images of monarch butterflies that can be used by other researchers to create automated approaches to count butterflies. **3)** We also create an open source tool that can be readily used by forest rangers, conservationists, and ecologists to estimate the counts of monarch butterflies. To the best of our knowledge, this paper creates the first open-source dataset consisting of annotated images of monarch butterflies as well as the first open source automated counting tool for the conservation of the species.

## 2 Preliminaries

Images: An input image *X* can be defined as a matrix of dimensions *m* × *n* × *k*, where *m* and *n* denote the height and the width of the image respectively, and *k* denotes the number of channels of the image. For example, a typical image in the “Joint Photographic Experts Group” format (jpeg) has three channels (*k* = 3), namely red, blue, and green. Each channel therefore consists of a matrix of size *m* × *n*. Each entry in an image is called a *pixel*, which refers to the smallest addressable element in the image. We use *x_ijk_* ∈ℤ^0+^ to denote the pixel in the *i*th row, *j*th column, and *k*th channel. In practice, *x* ∈ [0, 255]; therefore, each value *x_ijk_* can be represented by 8 bits (one byte) of memory. For all channels in an image in jpeg format, *x_ij_* = 0 represents the darkest version of the color and *x_ij_* = 255 represents the brightest version of the color. Fig. 1 illustrates how an input jpeg image is composed of three (red, green, and blue) channels.

**Figure 1:**
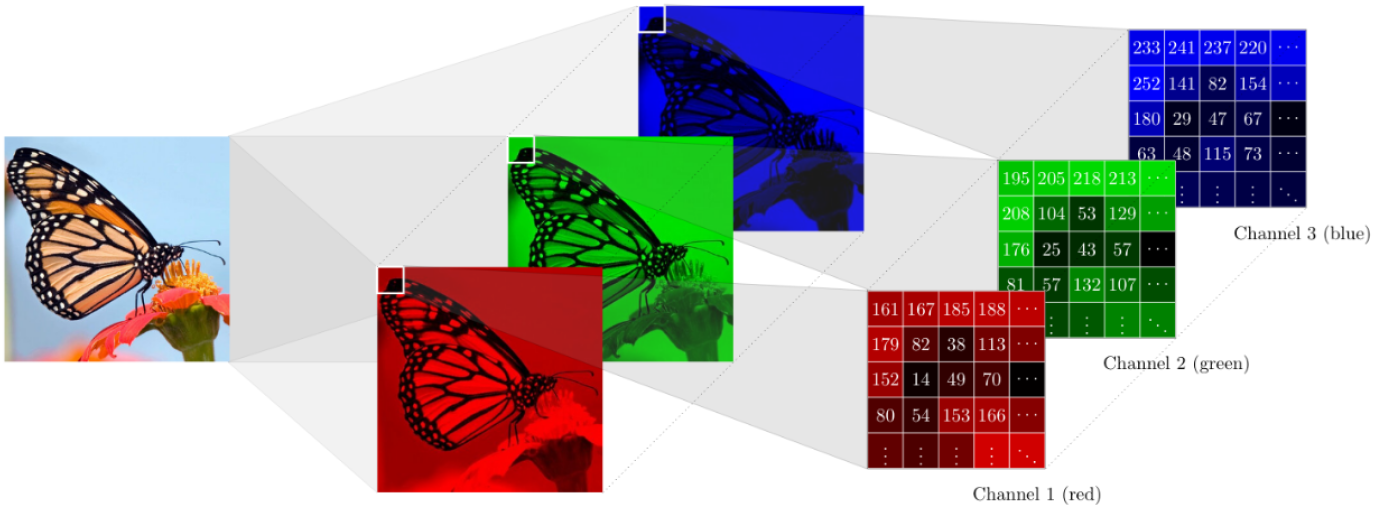
A sample image of a monarch butterfly. Images usually have multiple channels. Each channel can be represented by a matrix of non-negative integers

### Convolution

In image processing, convolution is defined as an operator that is used to multiply two matrices of the same dimensionality to produce a new matrix of the same dimension. While this is a rather loose definition, it is particularly well-suited to define the process of convolution. Usually, the convolution operator involves an element-wise multiplication of the matrices. As an example, consider two matrices *A* and *B* such that

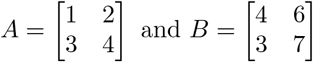

The convolution of *A* and *B*, denoted by *A* ∗ *B* creates a new matrix *D* (say) such that 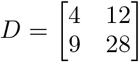. The idea of convolution is widely used in image processing to *smooth* images. To achieve this, a matrix *C* (say) is used which is typically much smaller than the original image. The smaller matrix is commonly referred to as a *filter* or a *kernel*. The matrix *C* is then slid through the original image matrix in a way that it moves through all positions where it fits entirely within the boundaries of the image [Fisher et al., 1996]. For each such position, the resulting value of the convolution is calculated in two steps. We use a single channel image *X* of dimensions *m* × *n* to illustrate the idea. Let the kernel matrix *C* be of size *l* × *l*. Let the portion of the original image that is obtained by placing the top-left corner of the matrix on pixel *x_ij_* be denoted by 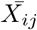. First, the convolution 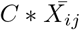 is calculated. This produces an intermediate matrix. The elements of the intermediate matrix are then added to create the corresponding output pixel. The overall operation results in an output image *Y* such that 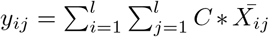. An example of the convolution operator is shown in fig. 2. A common method of choosing the kernel matrix is to sample it from an isotropic bivariate gaussian distribution. In theory, samples from such a distribution are non-zero across the two-dimensional space. In practice, however, it suffices to truncate the sample based on the required size of the kernel.

**Figure 2:**
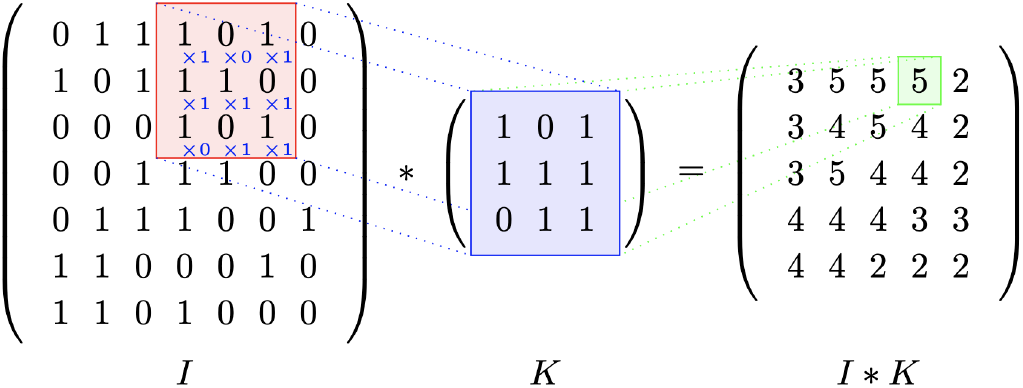
An example showing the convolution of a matrix *I* with a kernel *K*

### Neural Networks

Neural networks are computational models that use multiple processing layers to learn abstract representations of data [LeCun et al., 2015]. One of the most important computational models ever created, these methods have had widespread impact in various fields of research such as speech recognition, image recognition, image generation, and object detection. Neural networks consist of an input layer, an output layer, and multiple hidden layers. Each layer is composed of various *neurons* that represent computational nodes. Each neuron receives an input from the previous layer and calculates an output based on an *activation* function and *weights*. An intuitive way to think about the weights is to consider them as adjustable knobs [LeCun et al., 2015]. The output layer is the final layer of the network and it provides the output of the model. The goal of the learning procedure is then to optimize the knobs to minimize some pre-defined loss function. In practice, neural networks are optimized by gradient-based optimization approaches. A convolutional neural network (CNN) is a type of neural networks that are especially suited to working with images. CNNs typically consist of a convolution layer, in which a kernel of weights is slid across the image to calculate an element-wise product, which is then integrated across the channels of the image [O’Shea and Nash, 2015].

## 3 Approach

Consider a dataset *D* consisting of *q* images. We refer to the image matrix of the *p*th image by *X_p_*. Let the dimension of *X_p_* be *m* × *n* × *k* ∀*p* ∈ {1, … , *q*}, where *m*, *n*, and *k* denote the height, width, and the number of channels of the image respectively. For each image *X_p_*, let *Y_p_* denote a matrix of dimensions *m×n* that represents the annotated locations of butterflies in the image. We assume that each butterfly’s location can be represented by one pixel. This assumption does not affect the performance of our approach since the neighborhood of the annotated pixel plays a crucial role in the overall learning, as we show later. We consider that 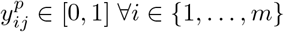 and ∀*j* ∈ {1, … , *n*}, where 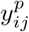 denotes the entry in the matrix *Y_p_* at the *i*th row and the *j*th column. Further, 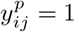 if and only if 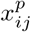 has a positive annotation for the presence of a butterfly. Therefore, the total number of butterflies in an image *X^p^* can be denoted by 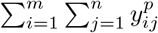.

Our goal is to predict the number of monarch butterflies in an arbitrary image. For an unseen image (an image unseen by our trained model) indexed by *r* (say), the image matrix *X_r_* is known. The annotations *Y_r_* are unknown. Our goal is to utilize the annotated dataset *D* learn a function that can predict 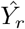 given an arbitrary *X_r_*. Formally, we want to learn *f* (*Y* | *X*), which denotes the conditional distribution of *Y* given *X*. Naturally, we want to minimize a loss function 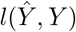 that measures the deviation or difference between the ground truth output and the predicted output. Instead of directly using *Y* to learn the function, it is common to create an intermediate density map of *Y* [Xie et al., 2018, Boominathan et al., 2016]. The creation of the density map alleviates the problem of detection and segmentation of individual objects in the image. The density map can be obtained by convoluting a gaussian kernel over the matrix *Y*. If the kernel is normalized to sum to one, then integrating over the density map provides the number of annotations in the original image. We illustrate this in Fig. 3.

**Figure 3:**
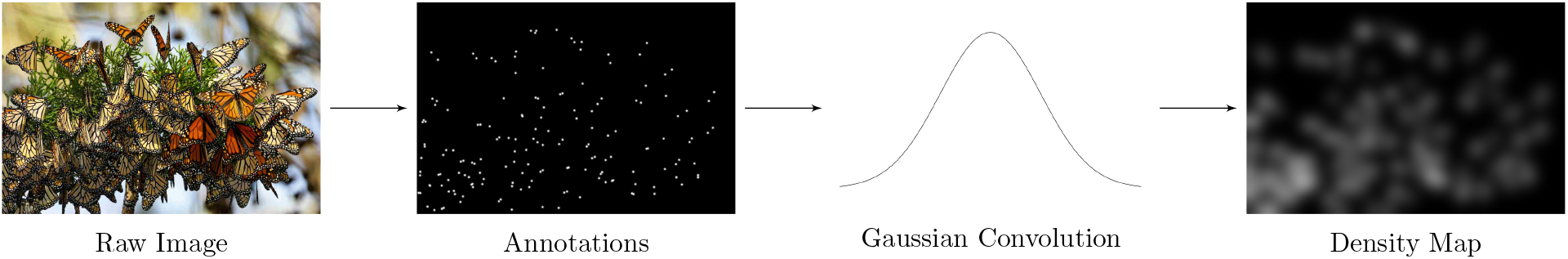
Butterflies are annotated manually. The resulting annotations are passed through a Gaussian convolution, which generates the final density map.

We use a Convolutional Neural Network (CNN) to learn the function *f* . Specifically, we use the U-Net architecture [Xie et al., 2018], which is a widely used architecture for image segmentation. The U-Net has a structure that represents an autoencoder [Goodfellow et al., 2016]. Given an input image, the key idea behind the U-Net architecture is to first process the image by a set of convolutional layers, followed by layers that downsample the image. This procedure is repeated several times before upsampling the resulting structure to match the size of the input image. The corresponding upsampling and downsampling units are coupled by concatenating the layers, due to which the architecture retains the essential features from the original image. The architecture of the U-Net used to predict monarch butterflies in a given input image is given in the Fig. 4.

**Figure 4:**
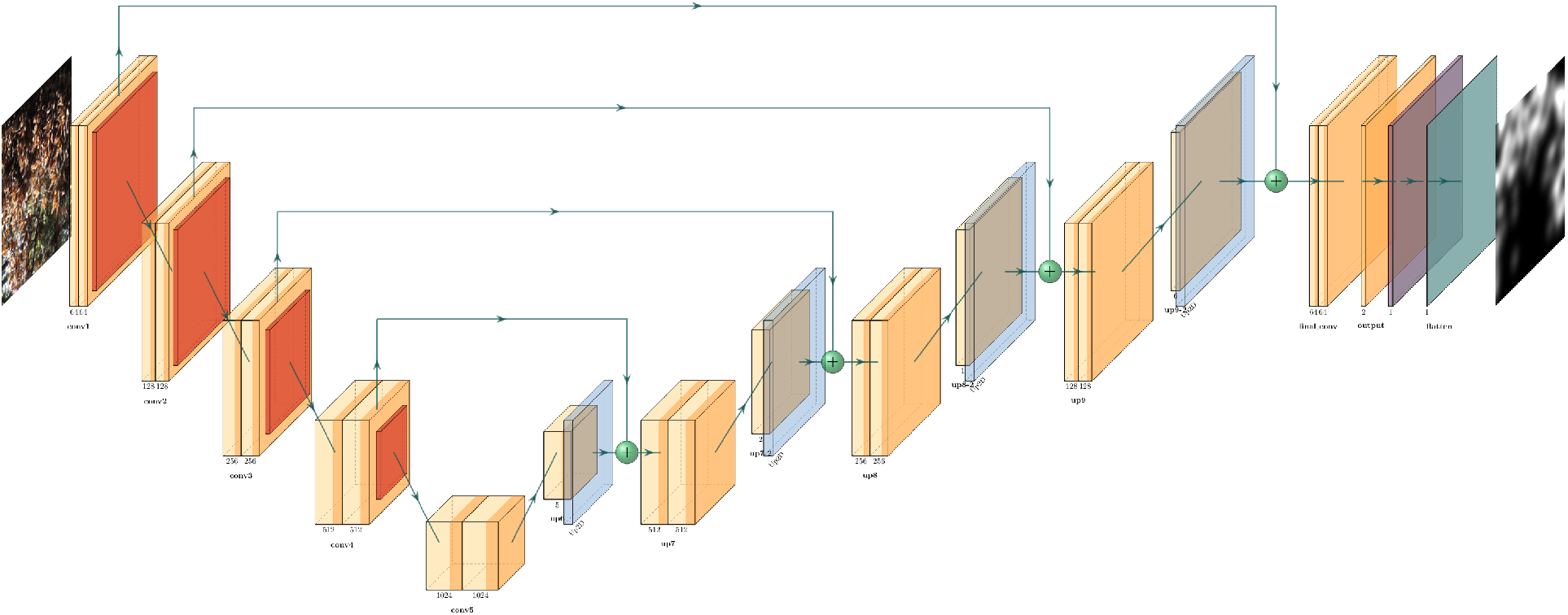
U-Net architecture. The network produces a density map for the given input image containing monarch butterflies.

An important step before using a CNN to learn the function *f* is to create accurate annotations. One needs to choose the location of the annotation carefully depending on the specific domain of application. For example, in images that contain dense crowd of humans, it is common to annotate the head of each person [Boominathan et al., 2016]. For monarch butterflies, we choose to annotate each butterfly on the upper most tip of one of their wings as shown in Fig. 5. The choice of location for annotation is based on the structure of clusters in which butterflies exist. Typically, monarch butterflies rest in dense clusters in wintering sites. The high density creates occlusion; as a result, oftentimes, the outer edges of their wings are the only parts visible and can be relied on for distinguishing butterflies. If the upper most point on a monarch butterfly’s wings was occluded by other butterflies in the cluster, any visible point on the butterfly’s body was marked.

**Figure 5:**
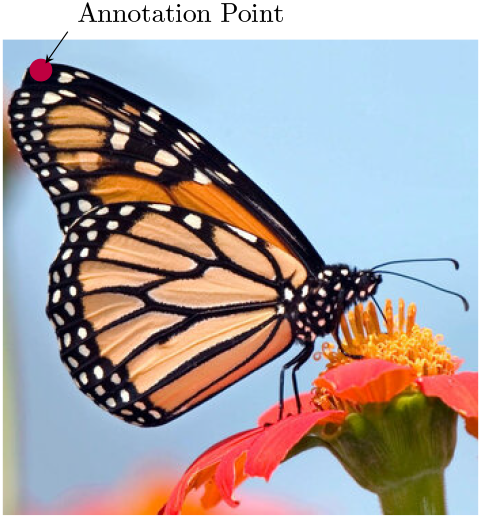
Where human annotators place labels on butterflies. When the upper wing tip of the butterfly is not visible, the annotations are placed on any part of the butterfly that is visible.

## 4 Data

Given the lack of open-source data on monarch butterfly clusters, we curated a dataset for this project. Monarch images were gathered by scraping the web. We collected 51 high-resolution images of varying sizes. We selected images primarily based on size and qualitative metrics like the existence of dense clusters and proper lighting. We avoided images of monarchs in flight since in practice, it is difficult to scan an area in the forest when monarch butterflies are in flight. We annotated each image using the online tool https://www.makesense.ai [Skalski, 2019]. The set of annotated images is available for the research community to use, and can be found at https://github.com/monarch-counter/data-public.

After all the monarch butterflies were annotated, the dataset was augmented by rotating and cropping varied sections of the images. We used a sliding window of size 1024 × 1024, and iteratively moved the window horizontally as well as vertically to crop the original images. The resulting images were then rotated about their centers for 0°, 90°, 180°, and 270°. This resulted in a total of 2,810 annotated images. We used a bivariate gaussian kernel with a mean of 1.0 and a constant standard deviation of 40.0 along both axes.

## 5 Experiments

### Setup

The processed raw images and derived density maps were split into a training set of 2308 images and a test set of 462 images. The model was trained on a Nvidia RTX 3080, a Graphics Processing Unit (GPU) with 10GB GDDR5 RAM and 8704 CUDA cores (34 TFLOPS). We use accuracy on unseen data (test set) as the primary metric for evaluation. For an input image *X_i_*, we define accuracy 100 - err_*i*_, where 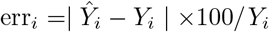. The predicted count of the output images is computed by integrating the output density generated by the neural network. The actual count of the images is retrieved from the ground truth annotations.

### Results

The mean error 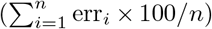 for all the test images (a total of *n* images) is 5.09%, resulting in an average accuracy of 95.91%. While there are no comparable baselines, we believe that the proposed approach, which is almost 96% accurate on test data, can be used in the field to estimate the count of monarch butterflies. We also analyzed the images to understand the results better. The distribution of errors is shown in Fig. 6a. An interesting observation is that the model identified butterflies that were missed by human annotators. We show this result in Fig. 7. Annotators can incorrectly label images due to several reasons. In the example shown in Fig. 7, the annotaters did not label butterflies in the background that are blurred and the label for the distinctly visible butterfly at the bottom of the image was cropped during pre-processing. However, the trained model, which learned abstractions using a large number of labeled images, predicted 4.87 butterflies in this case. This observation shows, to some extent, that the model is robust enough to decipher butterflies in poor quality images.

**Figure 6:**
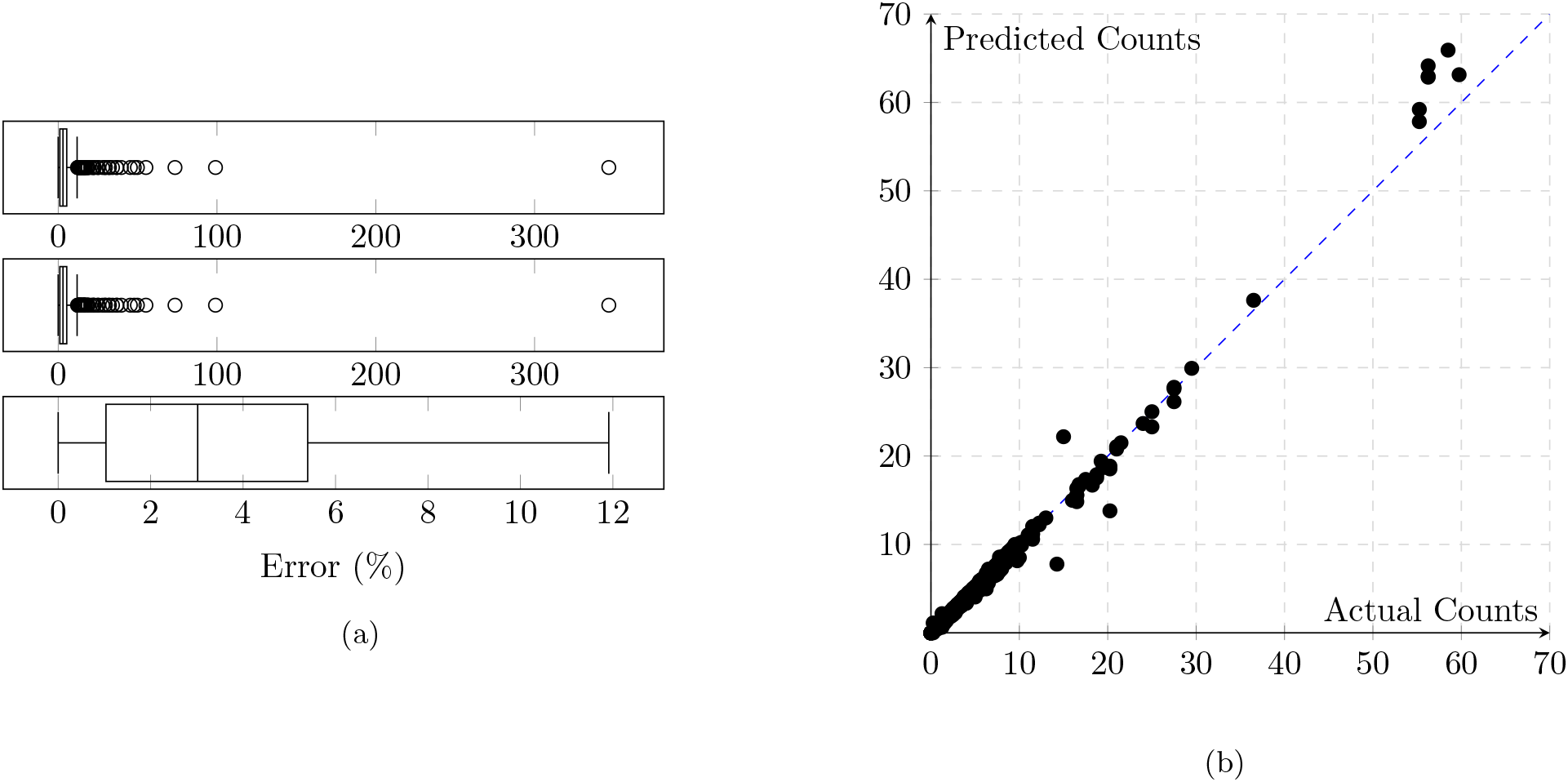
(**a**) Distribution of errors on unseen data. **Top**: Box plot of errors in images in the test set. **Middle**: Box plot of errors in images in the test set except for the furthest outlier. The furthest outlier is shown in Fig. 7, an image where human annotaters labeled a single butterfly but the model predicted 4.8 butterflies. **Bottom**: Box plot of errors in the test set images without outliers. (**b**) Predicted versus actual counts of monarch butterflies on unseen images (test set). Actual counts of images are annotated by humans. The blue dashed line represents *y* = *x* which denotes predictions with no error. We observe that the model based on deep neural network follows the diagonal line well.

**Figure 7:**
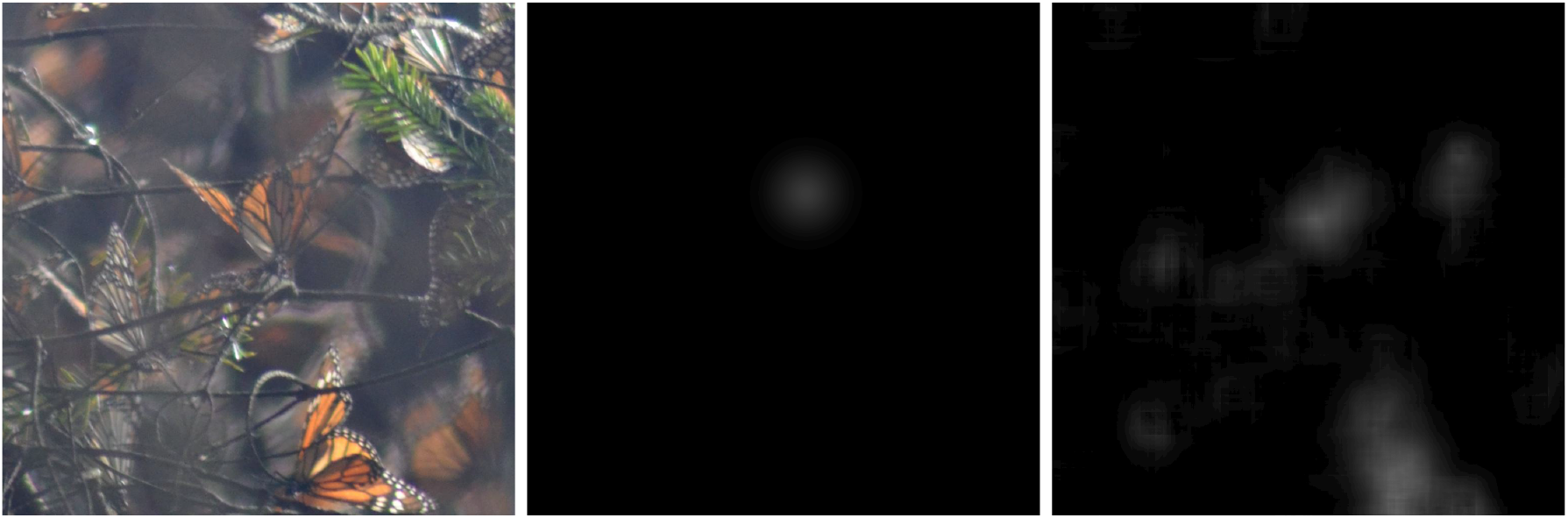
A sample image where annotations by humans missed butterflies in the background due to the blurred nature of the image. Also, the annotation for the distinctly visible butterfly was cropped during pre-processing. We find that the neural network is robust to such errors as it learns the necessary abstractions from a large number of images.

We also use the absolute error of prediction as a validation metric directly by comparing the difference between the expected and predicted butterfly counts. Fig. 6b shows how much a given test image’s predicted count varied from it’s expected count. On average, the model’s average error is 0.9, which means that it predicts 0.9 butterflies more (or less) than the ground-truth. For reference, the average number of butterflies per image in our test data is 20. We specifically analyzed the outliers in this case as well. As an example, we present the raw test image, expected density map, and model generated density map for the image in the test set with the largest absolute error in Fig. 8. Comparing the predicted density map to the raw input test image and expected density map indicates the model over-predicts butteflies in the extremely densely populated area of the image (the bottom part). We hypothesize that as more images are available from multiple annotators, the model’s performance in images such as these will improve.

**Figure 8:**
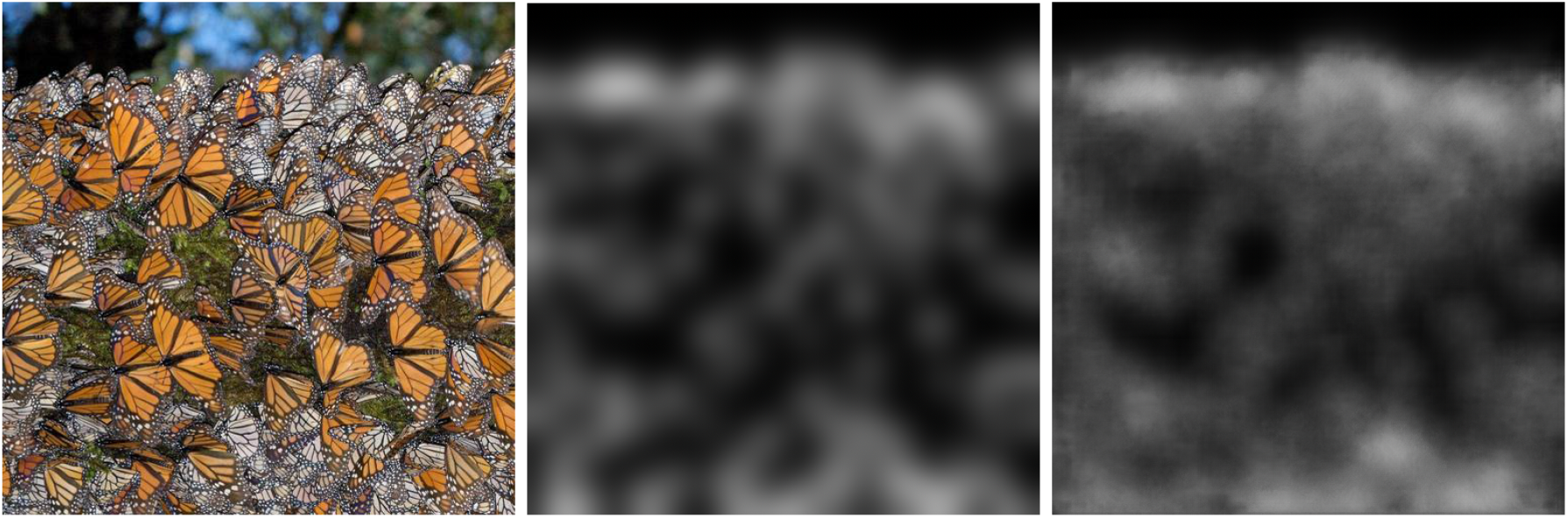
Image in the test set where our model performed the worst with respect to absolute error. From left to right, we show the actual image, the ground-truth density, and the predicted density. The ground-truth annotation (by humans) consisted of 225 butterflies, whereas the model predicted the presence of 256 butterflies. We hypothesize that the model predicts more butterflies due to parts of the image being extremely densely occupied by monarchs.

## 6 Open-source Implementation

In order to make this work easily accessible to practitioners, we release our codebase in two ways. An overview of our toolchain is shown in Fig. 9. The design choices for the tools were based on factors like ease of use, primary user demographic, and resource constraints. We describe the tools below.

**Figure 9:**
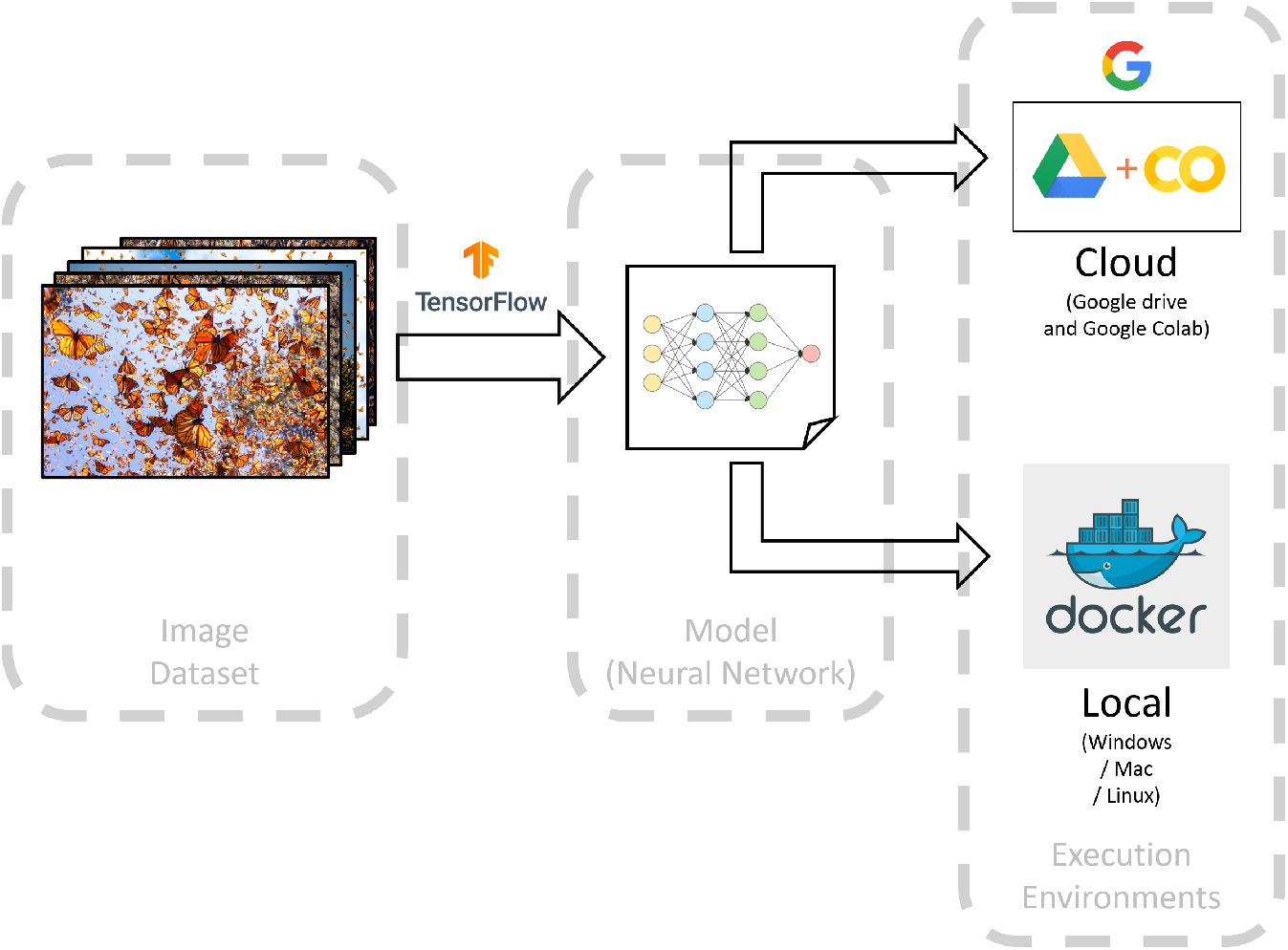
Overview of the tools used for developing the user interfaces

### Google Colab

[Bisong, 2019] is a cloud-based collaborative platform for prototyping machine learning models. It uses sophisticated hardware such as GPUs and TPUs located in Google’s cloud infrastructure for training machine learning models, which removes the overhead of acquiring the specialized hardware from the end user. Colab provides the end user with a web-based interface for accessing this hardware via a Jupyter notebook [Kluyver et al., 2016], which can be used to run Python scripts on the remote hardware in the cloud. These notebooks are self-contained environments that can be shared and worked on collaboratively. Our entire toolchain, in the form of an interactive Jupyter notebook on Google Colab platform can be found at https://tinyurl.com/y2vrhb97. The script in the notebook downloads pre-trained neural network model. The user can then upload one or more images of monarch butterflies to estimate counts in the images. The script runs the model on Google Colab and presents the results. The user is given the option to download the results in the form of a CSV (comma-separated values) file.

### Docker

[Merkel, 2014] is a software platform that helps developers package their code. By doing so, it makes the software application easier to ship and deploy. These packages are called *containers*, which contain the software application along with all its software dependencies. Creating containers helps the end user as the containers can be easily installed in the target system by using a simple set of commands. The model used to run and analyze the test results is packaged into a docker-cli (command line interface) and open-sourced for other researchers to use. The CLI takes the folder path of a folder filled with 1024px × 1024px images in jpeg format. It outputs the filename and count of each image in the folder on the console. The latest docker CLI image can be found via the tag `shruti222patel/monarch-counter:latest`.

## 7 Conclusion

In this paper, we present an approach to automatically count individual monarchs in dense clusters using deep learning. Further, we create a dataset consisting of over 2000 annotated images of monarch butterflies. The data as well as our implementation is open-source and packaged in the form of easy-to-use tools. We foresee that this tool can be used in two important ways. First, for the large overwintering population and dense clusters in Mexico, the method could be used to estimate the number of monarchs in selected clusters, and to extrapolate to estimate the number of monarchs in an overwintering colony. This could further help in determining the relationship between occupied hectares and monarch numbers [Thogmartin et al., 2017a], and provide a more accurate way to monitor monarch butterflies over time. Second, the tool could be used to quantify the number of monarchs in overwintering sites in California. While most North American monarchs overwinter in Central Mexico, many monarchs that live west of the Rocky Mountains fly shorter distances to the California coast to overwinter in groves of Eucalyptus and conifer trees [Urquhart and Urquhart, 1977, Yang et al., 2015, James et al., 2018]. Overwintering densities in these sites are much lower than those in Mexico [Malcolm, 2018], and as such, our method would be straightforward to implement to obtain accurate estimates of the entire monarch population sizes in this area. This would be especially important, because monarch densities in western North America have declined more precipitously than in Mexico [Schultz et al., 2017]. Our pre-trained model can also be used by researchers to aid conservation of other species. This is because even though our model shows impressive performance, we are currently investigating its robustness against noise in the annotation process.

## Author Credits and Affiliations

**SP** (Independent Researcher): Methodology, Software (Machine Learning, Tool Development), Investigation, Writing (Original Draft), Project Administration: **AK** (Vanderbilt University): Methodology, Software (Data Processing, Machine Learning, Tool Development), Investigation, Data Curation, Writing (Review and Editing), Computing Resources; **AM** (Vanderbilt University): Conceptualization, Methodology, Software (Data Processing, Machine Learning), Investigation, Writing (Original Draft, Review and Editing), Data Curation, Supervision, Project Administration; **KG** (Middle Tennessee State University): Data Curation, Software (Data Processing, Tool Development); **JDR,** (Emory Univeristy): Writing (Review and Editing), Supervision.

## Notes

https://github.com/monarch-counter

### Competing Interest Statement

The authors have declared no competing interest.

### Summary of Updates

Author affiliations updated.

